# Elevated mutation rates in the multi-azole resistant *Aspergillus fumigatus* clade drives rapid evolution of antifungal resistance

**DOI:** 10.1101/2023.12.05.570068

**Authors:** Michael J. Bottery, Norman van Rhijn, Harry Chown, Johanna L. Rhodes, Brandi N. Celia-Sanchez, Marin T. Brewer, Michelle Momany, Matthew C. Fisher, Christopher G. Knight, Michael J. Bromley

**Affiliations:** Manchester Fungal Infection Group, Division of Evolution, Infection, and Genomics, Faculty of Biology, Medicine and Health, University of Manchester, Manchester, United Kingdom; Department of Medical Microbiology, Radboud University Medical Centre, Nijmegen, Netherlands; Fungal Biology Group and Department of Plant Biology, University of Georgia, Athens, GA 30602, USA; Medical Research Council Centre for Global Infectious Disease Analysis, Imperial College London, London, United Kingdom; Department of Earth and Environmental Sciences, School of Natural Sciences, Faculty of Science and Engineering, The University of Manchester, United Kingdom

## Abstract

The evolution of antifungal resistance is an emerging global threat. Particularly concerning is the widespread occurrence of azole resistance within *Aspergillus fumigatus*, a globally ubiquitous environmental mould that causes over 1 million life-threatening invasive infections in humans each year. It is increasingly evident that the environmental use of azoles has led to selective sweeps across multiple genomic loci resulting in the rapid expansion of a genetically distinct cluster of genotypes (clade A) that results in resistance to clinically deployed azoles. Isolates within this cluster are more likely to be cross resistant to agricultural antifungals with unrelated modes of action suggesting they may be adapting rapidly to antifungal challenge. Here we show that this cluster is not only multi-azole resistant but has increased propensity to develop resistance to new antifungals because of variants in the DNA mismatch repair system. A variant in *msh6* is found almost exclusively within clade A, occurs in 88% of multi-azole resistant isolates harbouring the canonical *cyp51A* azole resistance allelic variant TR_34_/L98H, and is globally distributed. Naturally occurring isolates with this *msh6* variant display a 4 to 9-times higher rate of mutation, leading to an increased propensity to evolve resistance to current and next generation antifungals. We argue that pervasive environmental use of fungicides creates selective arenas whereby genotypes of *A. fumigatus* with increased adaptive capability thrive in the face of strong directional selection, leading to the genesis and amplification of antifungal resistance. These results help explain the pronounced clustering of multiple independent resistance mechanisms within the mutable clade A. Our findings further suggest that resistance to next generation antifungals is more likely to emerge within organisms that are already multi-azole resistant, posing a major problem due to the prospect of dual use of novel antifungals in clinical and agricultural settings.

## Main

*Aspergillus fumigatus* is a globally prevalent saprotrophic mould that causes a wide spectrum of diseases in humans. Globally 30 million people are at risk of contracting a life-threatening invasive aspergillosis (IA) infection due to the impact of immunosuppressive medications, neutropenia, and comorbidities such as COPD (*1*). Over 1 million people develop IA annually (*2*, *3*), with mortality rates ranging between 30 to 60% even where gold standard treatments are given (*4*). In addition, 3 million people are estimated to have chronic infections caused by *Aspergillus* spp. (chronic pulmonary aspergillosis; CPA) with around 10% succumbing to their infection in the first year and 5-year survival rates of 62%. With a growing at-risk population, these infections are of critical concern resulting in *A. fumigatus* being listed by the WHO as a critical priority fungal pathogen (*5*, *6*).

Resistance to azoles, the primary class of antifungal for the treatment of aspergillosis, is a growing problem with prevalence reported as high as 10% in some centres (*7–9*). Mortality rates in those infected with resistant isolates increases by 25% even if alternative salvage therapies are provided (*10–13*). The evolution of azole resistance during treatment of patients with chronic disease is typically associated with the acquisition of specific *de novo* non-synonymous mutations in *cyp51A* (*14*), leading to changes in the target of azoles – sterol-14α-demethylase – which catalyses a critical step in the synthesis of ergosterol, an essential component of the fungal cell membrane. There is also compelling evidence that the widespread use of agricultural fungicides functionally analogous to clinical azoles select for pan-azole resistance in the environment resulting in a hallmark set of mutations in *cyp51A* (*15*). Environmental pan-azole resistance is characterised by a 34-base pair tandem repeat in the promoter of *cyp51A* which leads to the over-expression of the gene, coupled with point mutations in *cyp51A* (notably the substitution of leucine 98 with histidine) which reduce the binding affinity of the azoles (*16*). The first isolates with TR_34_/L98H (*17*), which provide high levels of resistance to itraconazole and voriconazole, were observed in the Netherlands in both environmental and clinical isolates, prompting the hypothesis that environmental selection is driving resistance (*18*). Subsequently, isolates with these mutations and similar sets of mutations (e.g. TR_46_/Y121F/T289A(*19*)) have been identified globally (*15*). Genome sequencing and molecular epidemiology has now identified multiple examples of near identical genotypes from both environmental and clinical sources (*20*) providing strong evidence for patient acquisition of azole-resistant *A. fumigatus* from environmental sources.

Despite the fact that *A. fumigatus* can undergo a sexual cycle and can also readily disperse on air currents due to the production of aerosolised conidia during asexual reproduction, recent genomic studies have identified strong population structuring linked to the presence of pan-azole resistance mutations (*20*, *21*). Phylogenetic analyses of *A. fumigatus* isolates from the United Kingdom showed pronounced genetic clustering into two populations, clade A and clade B, with the azole resistance mutation TR_34_/L98H almost uniquely occurring in clade A and wild-type *cyp51A* predominantly occurring in clade B (*20*). A third divergent azole-sensitive clade has also been identified within global isolate collections (*21*). This population genetic structure is reinforced by low levels of recombination between clades and high rates of recombination within clades (*21*). Moreover, the genomes in clade A show the signatures of strong directional selection across multiple loci known to be involved in resistance to clinical and agricultural fungicides (*20*). Intriguingly several other variant loci, not currently associated with resistance to agricultural fungicides are also enriched in this clade. Whether these loci have become enriched through hitchhiking with beneficial resistance mutations which are undergoing a selective sweep or are playing a direct role in the evolution of the clade remains unclear. The near uniform association between cross resistance across antifungal classes (*22*) and strains in clade A led us to hypothesise that this clade was evolving more rapidly in response to antifungal challenge, perhaps driven by the impact of these variants.

Defects in DNA mismatch repair (MMR) can result in elevated mutation rates in bacterial, fungal and mammalian cells (*23*). Elevated mutation rates in mutator microorganisms can confer an evolutionary advantage when adapting to novel, stressful or fluctuating environments, but are often associated with a fitness cost in stable environments due to the accumulation of deleterious mutations. The frequency of mutator strains within a population can be rapidly amplified through genetic hitchhiking upon beneficial adaptive mutations and can play an important role in the evolution of pathogens to novel stresses (*24–26*). Mutator phenotypes are key drivers of within-host evolution in response to the host immune system, abiotic stress, and antibiotic treatment, particularly during long-term chronic infection. The selection for elevated mutation rates has been identified in human fungal pathogens including *Candida albicans* (*27*)*, Candida glabrata* and *Cryptococcus neoformans* (*29*, *30*), and may play a role in the acquisition of resistance in *A. fumigatus* (*31*). But critically, elevated mutation rates are likely to influence the ability to adapt to strong fluctuating directional selection within the environment for example through the use of multiple different fungicides which will exert selection for genomic flexibility. We therefore investigated if polymorphisms in the MMR system were enriched in the multi-azole resistant clade A.

### Variants in mismatch repair genes are overrepresented in clade A isolates with TR34/L98H

Eukaryotic MMR consists of two major recognition complexes: MutSɑ (Msh2-Msh6), which recognizes base-base mismatches and small loops, and MutSβ (Msh2-Msh3) which recognises larger loops, with a bias towards deletion loops (*32*). The MutLɑ protein complex (Pms1-Mlh1) directs downstream protein-protein interactions, is required for daughter strand recognition, and has endonuclease activity. We screened 218 previously sequenced *A. fumigatus* isolates (*20*) originating from environmental (65) and clinical (153) sources in the United Kingdom, of which 91 contain TR_34_/L98H and 7 contain TR_46_/Y121F/T289A *cyp51A* azole resistance mutations, for variants in MMR genes *msh2* (AFUB_039320, AFUA_3G09850), *msh3* (AFUB_090020, AFUA_7G04480), *msh6* (AFUB_065410, AFUA_4G08300), *pms1* (AFUB_029050, AFUA_2G13410) and *mlh1* (AFUB_059270, AFUA_5G11700). In total, across all five genes 212 non-synonymous point mutations were present relative to the *Af*293 reference strain (Supplementary Table 1). No loss of function or truncation mutations were observed in any of the isolates, thus variants were expected to alter rather than abolish the activity of the MMR systems. Of these variants, non-synonymous mutations in *msh2* (c.A2435G, p.E812G), *msh6* (c.G698C, p.G233A) and *pms1* (c.A1331G,p.E444G) were significantly associated with clade A (Figure 1a,b,c, *msh2*: d.f. = 1, χ^2^ = 8.88, P = 0.0029, *msh6*: d.f. = 1 χ^2^ = 141.64, P < 2.2×10^-16^, *pms1*: d.f. = 1, χ^2^ = 19.12, P = 1.2×10^-5^), however, only the G233A variant in *msh6*, which occurs prior to the annotated N-terminal MutS domain responsible for mismatch recognition (Figure 1e), was significantly associated with the presence of the azole resistance mutation TR_34_/L98H in *cyp51A* (d.f. = 1, χ^2^ = 122.27, P < 2.2×10^-16^). With the exception of the variants we have identified within clade A of *A. fumigatus*, the G233 amino acid is perfectly conserved across over 150 million years of evolutionary history (*33*) within the Trichocomaceae family (Figure 1f). In total, 85% (105/123) of clade A isolates contained the G233A variant allele of *msh6*, while the variant is only present in 3% of isolates in clade B (3/95). Of the 86 TR_34_/L98H azole resistant genotypes 96.5% contained G233A, whereas of the 7 isolates containing the TR_46_/Y121F/T289A resistance haplotype, only 4 harbour the G233A *msh6* variant (Figure 1d). Notably, *msh6* is encoded on chromosome 4 and closely linked with *cyp51A,* 0.36 Mbs away. The 10-kb region in which it is encoded has a high average fixation index (F*_ST_*) of 0.1995 between isolates within clade A and B implying population subdivision at this locus (average F*_ST_* = 0.1312, F*_ST_* of *cyp51A* = 0.119). In addition, previous analysis shows this region has a significant association with itraconazole resistance (*20*) (treeWAS P<0.001). The association between G233A, clade A and TR_34_/L98H was also evident in a global collection of isolates(*34*) (Figure S1, Clade A: d.f. = 2, χ^2^ = 463.93, P < 2.2×10^-16^, TR_34_/L98H: d.f. = 1, χ^2^ = 326.23, P < 2.2×10^-16^). Thus, the presence of the non-synonymous variant G233A in *msh6* which is an essential component of MutSɑ, responsible for recognising base-base mispairing, is strongly associated with the presence of azole resistance allele TR_34_/L98H in clade A.

**Figure 1.**
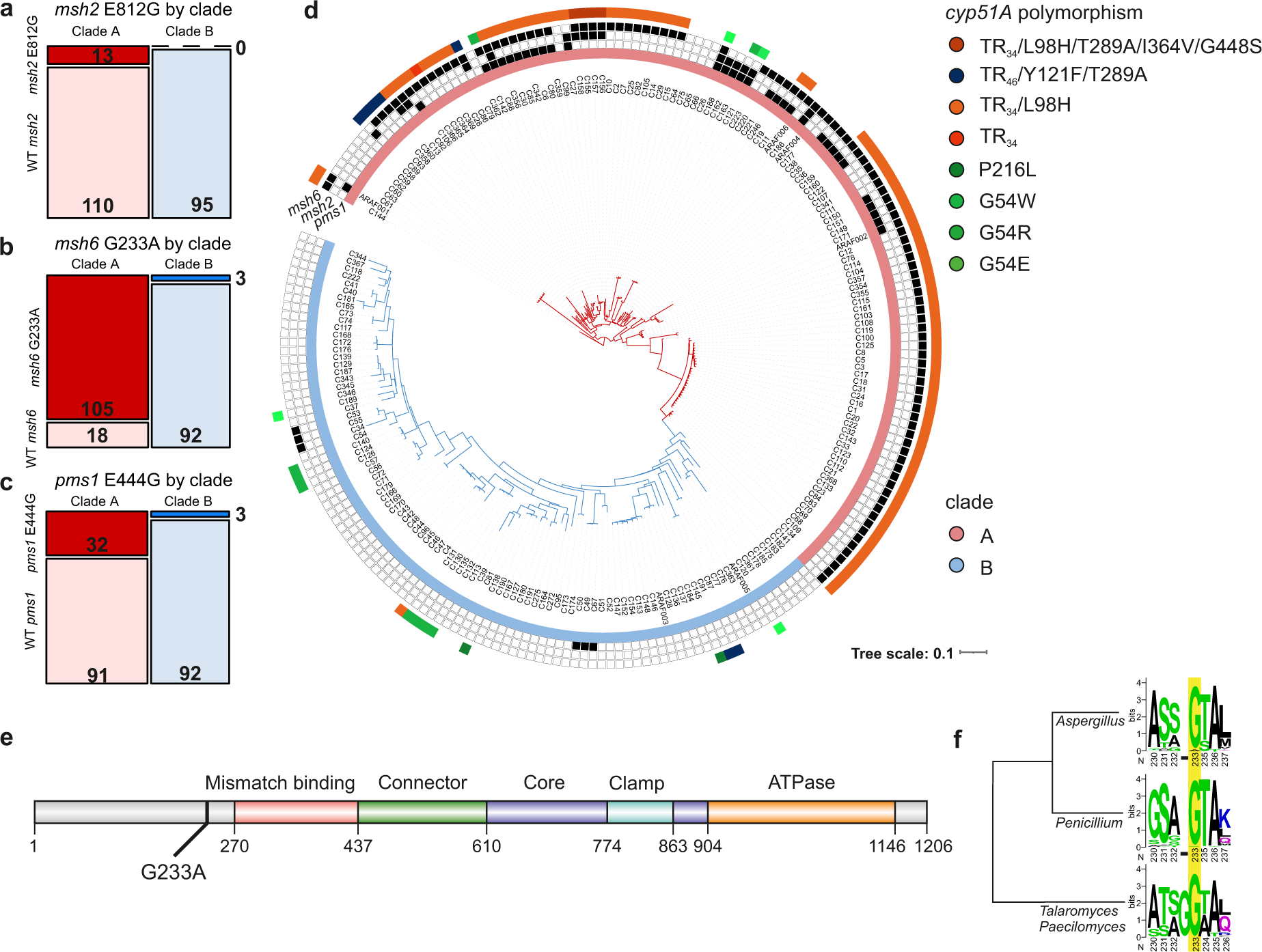
Variants in MMR are significantly overrepresented in clade. **A.** Occurrence of MMR variant alleles in clade A (red) and clade B (blue) for **a** *msh2* (E812G) **b** *msh6* (G233A) and **c** *pms1* (E444G). **d** An unrooted maximum likelihood phylogenetic tree using genome wide SNPs relative to *Af*293. The presence of variants in *msh2*, *msh6* and *pms1* are highlighted in the black boxes, the presence of *cyp51A* resistance variants and clade are coloured. **e** Domain structures of Msh6 in *A. fumigatus*, G233A variant labelled, labels show predicted domain positions in protein sequence. **f** G233 locus homology across Trichocomaceae, alignments of 140 isolates spanning *Aspergillus ssp.*, 27 *Talaromyces* and *Paecilomyces*, and 122 *Penicillium* Msh6 protein sequences. G233 highlighted in yellow, G233A variants are only present in *Aspergillus fumigatus*. Cladogram shows hierarchical clustering of Msh6 protein sequences.

### MutS and MutL null mutants result in a hypermutator phenotype

Since variant alleles in *msh2*, *msh6* and *pms1* are significantly associated with clade A, and in the case of *msh6* TR_34_/L98H azole resistance genotypes, we first asked whether each of the three genes influence mutation rate. The three genes were independently deleted from the WT strain MFIG001 (*35*). The minimal inhibitory concentrations to voriconazole, a current antifungal used to treat *A. fumigatus* infections, or the phase III clinical trial compound olorofim in the orotomide class (*36*) were not altered in the MMR defective strains relative to the parental MFIG001 strain (Figure S2) indicating no direct effect of these alleles on azole or orotomide sensitivity. To measure mutation rates a modified Luria-Delbrück fluctuation test (*37*) was implemented in which replicate cultures grown without selection were challenged with voriconazole to determine the probability that mononucleated spores would spontaneously gain mutations that provide resistance (Figure 2a). The rate of spontaneous mutation in the WT MFIG001 strain to voriconazole was 2.78⤬10^-10^ (± 6.9⤬10^-11^) per spore, similar to rates measured in other fungal species (*38–40*). The modified Luria-Delbrück method was validated by treatments with the mutagen ethyl methanesulfonate during growth. The method detected the linear increase in mutation rate in MFIG001 to voriconazole with increasing concentrations of the mutagen (Linear regression, R^2^ = 0.92, F_1,13_ = 153, P < 0.001, Figure S3). To detect changes in the mutation rates of MMR deletion strains they were tested with the modified Luria-Delbrück method. Mutation rates for voriconazole resistance in *Δmsh2*, *Δmsh6* and *Δpms1* were ∼80-, ∼42 and ∼177-fold higher than the parental wild-type strain (Figure 2b, Two sample ML-test, MFIG001-*Δmsh2* T = - 16.3, P < 0.0001, MFIG001-*Δmsh6* T = −14.49, P < 0.0001, MFIG001-*Δpms1* T = −12.889, P < 0.0001). The MICs of spontaneous resistant mutants to voriconazole ranged from 4 µg/ml to 32 µg/ml and were not dependent upon the genetic background (Kruskal-Wallis, d.f. = 2, χ^2^ = 4.25, P = 0.119). Sequencing of *cyp51A* showed that 37% (10/27) of the randomly selected spontaneously resistant isolates had the known voriconazole resistance allelic variant G448S, but these exclusively occurred within isolates with MICs ≥ 16 µg/ml. No mutations within *cyp51A* were observed in the other resistant isolates and tandem repeats in the promotor of *cyp51A* were never observed. The frequency of resistance to another azole-class antifungal, itraconazole and olorofim showed similar significant increases in mutation rate in all three MMR deficient strains, with the largest increases in *Δpms1* and the lowest increases in mutation rate in *Δmsh6* (Pairwise Wilcoxon tests, P < 0.05, Figure 2c, 2d). The probability of resistance arising differed between antifungals (ANOVA, F_2,225_ = 42.49, P < 0.001), with resistance arising between 2 and 15 times more frequently to itraconazole than olorofim or voriconazole (Tukey post hoc test, P < 0.001), however, there was no difference between the frequency of resistant mutants to voriconazole and olorofim (Tukey post hoc test, P = 0.828). Of the randomly selected olorofim spontaneous resistant mutants 26/27 had mutations in the *pyrE* resistance hotspot G119 (*41*) which provided high levels of olorofim resistance (>2 µg/ml). Together, these results show that *msh*2 and *pms1* null mutants which abolish the activity of the MMR system result in highly elevated mutation rates that facilitate the emergence of resistance. Moreover, although MutSβ recognition complexes remains functional when disrupting *msh6*(*32*), the loss of function of *msh6* still results in significant increases in mutation rate.

**Figure 2.**
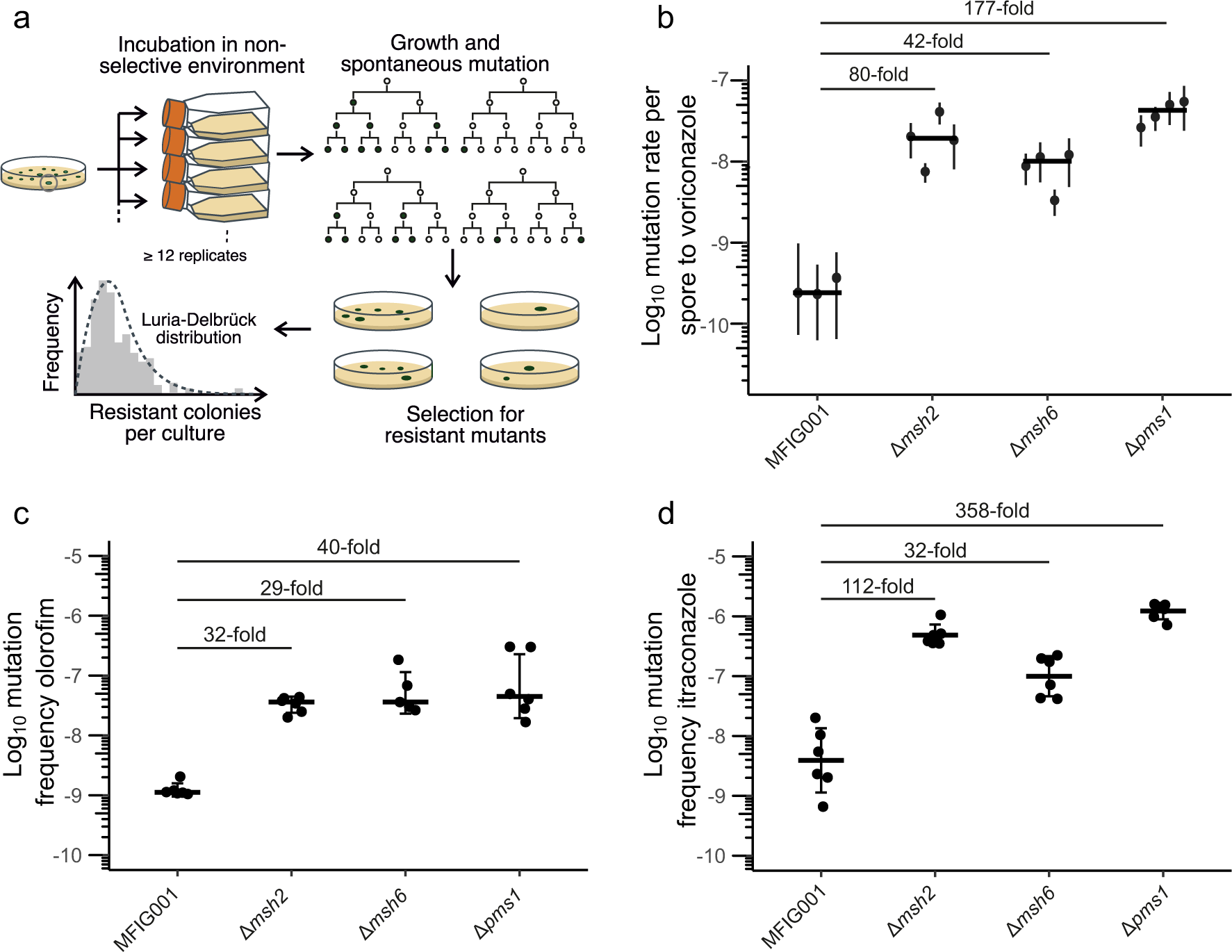
MutS and MutL null mutants significantly elevate mutation rates. **a** Workflow of fluctuation test to measure mutation rates in *A. fumigatus*. Clonal isolates were cultured in the absence of antifungal selection to generate genetic diversity. Resistant mutants were selected on lethal concentrations of antifungals. Counts of resistant mutants were fitted to the Luria-Delbrück distribution to calculate the number of mutational events. **b** Mutation rates for resistance to voriconazole for MMR deficient mutants. Each point shows the calculated mutation rate from a single independent fluctuation test using 12 replicate cultures. Error bars show 95% confidence intervals, cross bars show median mutation rate across fluctuation tests. Fold differences show median fold change in mutation rate from the parental MFIG001 strain. **c** Mutational frequency to olorofim resistance. **d** Mutational frequency to itraconazole resistance. Each point shows the mutational frequency of an individual population (N = 6). Error bars show SEM, cross bars show median mutational frequency. Fold differences show median fold change in mutation frequency from the parental MFIG001 strain.

### Defective MMR results in significant reduction in fitness

Uncontrolled mutation can result in the accumulation of deleterious mutations which decrease the fitness of hypermutator strains within stable environments (*42*, *43*). We therefore asked whether such costs were associated with the MMR defects in *A. fumigatus*. Though we detected non-synonymous variants in *msh2*, *msh6* and *pms1* no predicted loss of function mutations within the MMR genes were observed in the clinical or environmental isolates sequenced, suggesting that the complete loss of MMR is associated with a significant cost in *A. fumigatus*. However, no defect in radial growth rates were observed over 96 hours in the MMR deletion mutants relative to their wild-type parental strain in either nutrient rich or minimal growth conditions (Figure S4). Interestingly, morphological sectoring, likely to occur due to mutations during hyphal growth, occurred within the MMR mutants but not the wild-type. To determine whether fitness costs manifested over longer periods of growth, MMR deficient mutants were directly competed with the wild-type parental strain over five serial transfers on complete (rich) and minimal solid media (Figure 3). The frequency of MMR deleted strains decreased through time and was dependent upon both the MMR mutant and the environment (Mixed effects linear model, Transfer:Media F_1_ = 7.8447, P < 0.01, Transfer:Strain F_2_ = 7.8528, P < 0.001, Figure 3a). Growth media had a significant effect on the relative fitness in all the MMR deficient strains, with fitness being consistently lower in minimal media compared to rich media (Wilcoxon test, W = 233, P < 0.05. Figure 3b). Deletion of *msh2* resulted in the highest overall cost relative to the wild-type isolate, displaying 30% cost in rich media (T.test, T_5_ = −13.6, P < 0.001, Holm adjusted for multiple testing) and a 75% cost in minimal media (T.test, T_5_ = −30, P < 0.001, Holm adjusted for multiple testing, Figure 3b). Deletion of *pms1* also resulted in significant cost (Rich: 13% fitness cost, T.test, T_5_ = −4.9, P < 0.05, Minimal: 25% fitness cost, T.test, T_5_ = −15.5, P < 0.001, Holm adjusted for multiple testing), although lower than Δ*msh2* (Two sample t.test, T_9.68_ = −4.8, P < 0.0001). In contrast, the *msh6* null mutant did not have a significant decrease in relative fitness measured at transfer 5 when competed in either rich or minimal media (Figure 3), however the fraction of Δ*msh6* had reduced significantly by transfer 4 when competed in minimal media (Wilcoxon test, W = 0, P < 0.01). Thus, over longer periods of time there are significant costs associated with the loss of function of MutS or MutL complex.

**Figure 3.**
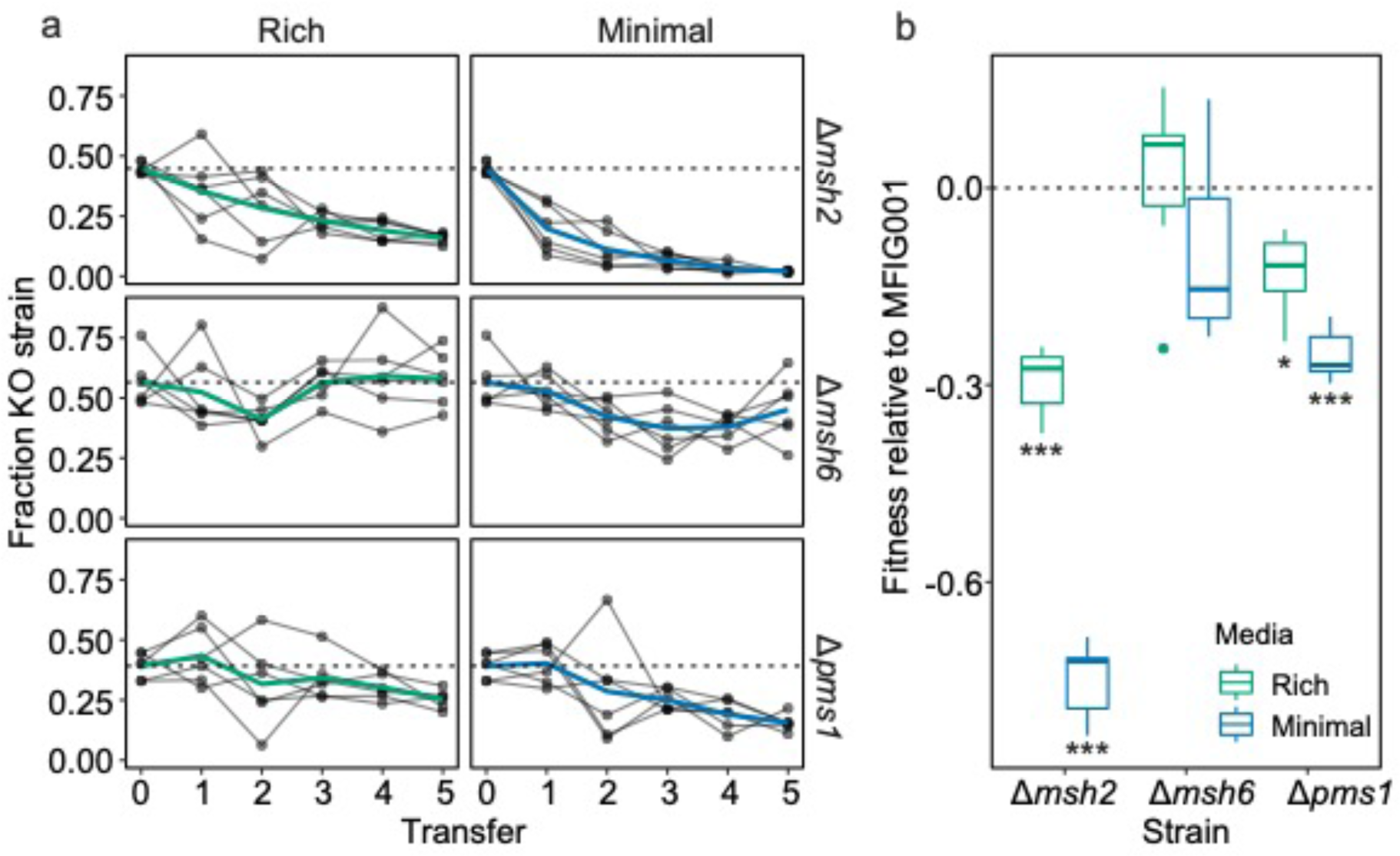
Deletion of MMR genes carries a significant cost to fitness. **a.** The fraction of *Δmsh2, Δmsh6* and *Δpms1* through time when in direct competition with the parental MFIG001 strain on solid agar faceted by media type (rich = aspergillus complete media, minimal = aspergillus minimal media). The coloured lines show the mean of 6 independent competitions, presented by individual grey lines. The horizontal dashed lines show starting fraction of KO strain. **b** Mean fitness of *Δmsh2, Δmsh6* and *Δpms1* relative to MFIG001 across five transfers, the cross bar shows the median (N=6), the lower and upper hinges correspond to the first and third quartiles and the whiskers extend to 1.5 * IQR. Box plots coloured by media type. The horizontal dashed line shows equal fitness, asterisks represent significant differences from zero (* < 0.05, ** < 0.01, *** < 0.0001).

### Msh6 G233A increases mutation rate and is correlated with increased mutation rates in clade A

Since we find deletion of MMR components to be costly, as well as increasing rates of anti-fungal resistance, we asked whether the more subtle variants we see in our strain collection have a similar effect. Therefore, we reconstructed the G233A variant of *msh6* within MFIG001 through marker-less CRISPR-Cas9 mediated transformation (*44*) to determine the effect of the variant on mutation rate. The *msh6* variant resulted in a modest but significant increase in rates of resistance mutation to olorofim of 3.6-fold relative to the isogenic parental strain in three independent transformants (Figure 4c, Two sample ML-test, MFIG001-*msh6*::G233A T = −2.9576, P < 0.01). We therefore hypothesise that this variant will be associated with variation in mutation rates in natural isolates. To test this hypothesis we assayed the rate of olorofim resistance mutation in 10 isolates with a range of variants in *msh2*, *msh6* and *pms1* from clade A and clade B (Table 1). Only the novel antifungal olorofim could be used for these fluctuation tests as the clade A isolates and one clade B isolate already had mutations in *cyp51A* that conferred resistance to azoles. The olorofim MICs of the isolates were not significantly different between isolates, and all fell at least 4-fold below the concentration used to select for resistant mutants (Figure S5) enabling direct comparisons of mutation rates to be made. The isolates from clade B without variants in the MMR genes did not have significantly different mutation rates compared to MFIG001, including a clade B isolate with *cyp51A* TR_46_/Y121F/T289A azole resistance mutations (Figure 4b, Two sample ML-test, MFIG001-clade B isolates, all P > 0.05). In contrast, isolates from clade A with the *msh6*::G233A variant had between a 3.5- and 7.4-fold increase in mutation rate (median 5.7-fold), significantly increasing the likelihood of spontaneous olorofim resistance arising (Figure 4d, Two sample ML-test, MFIG001-*msh6*::G240A isolates, all P < 0.05). These isolates encompassed both TR_34_/L98H and TR_46_/Y121F/T289 *cyp51A* resistance genotypes and isolates with clinical and environmental origins (Table 1). One clade A isolate, the only one without *msh6*::G233A, C89, did not display similarly elevated mutation rates. These results show that the presence of G233A variant in *msh6*, unique to clade A, is associated with elevated rates of mutation to resist a novel antifungal in natural isolates. Moreover, this elevated mutation rate was not influenced by the genotype of the linked of azole resistance locus.

**Figure 4.**
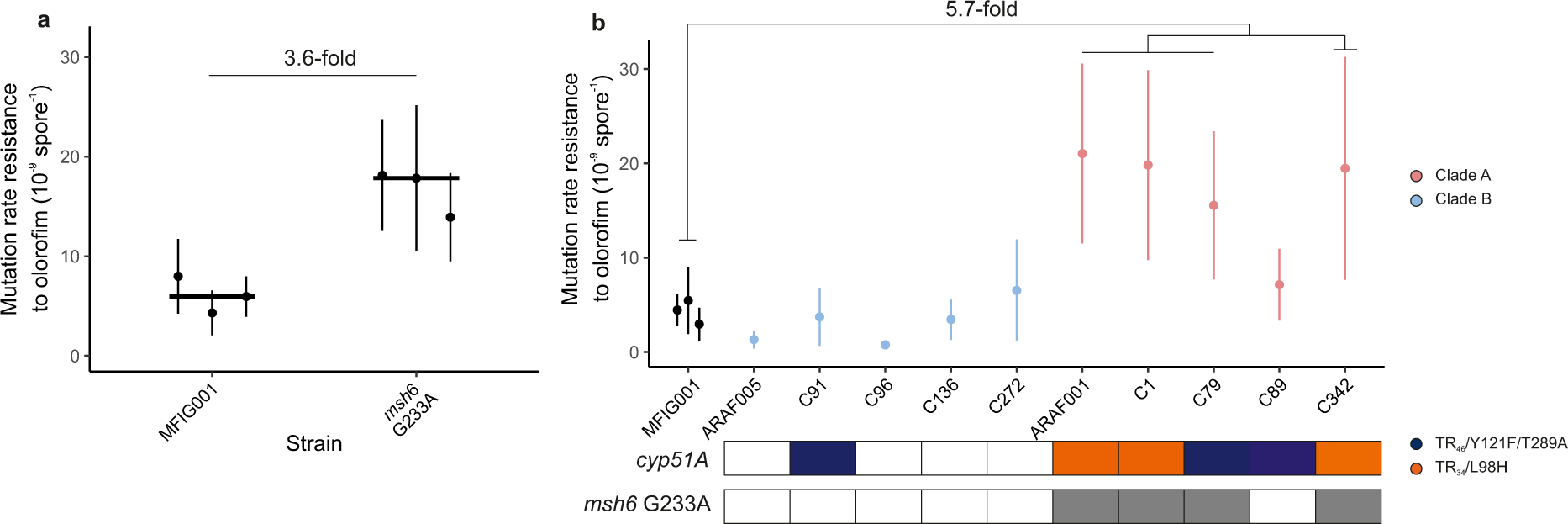
*msh6* G233A elevates mutation rates in neutral background and in natural isolates from clade A. **a** Each point shows the calculated mutation rate from a single independent fluctuation test using 12 replicate cultures. Each of the three points for the *msh6* G233A variant is a separate transformant. Error bars show 95% confidence intervals. Fold change shows the median fold change in mutation rate from the parental MFIG001 strain. **b** Mutation rate of natural genotypes to olorofim. Each point shows the calculated mutation rate from a single independent fluctuation test using 12 replicate cultures. Error bars show 95% confidence intervals. Fold difference show the median fold change in mutation rate of isolates with G233A allele from the parental MFIG001 strain. Points coloured by clade, red clade A, blue clade B, black MFIG001. Key below plot shows the presence of azole resistance mutation in *cyp51A*, and the presence of *msh6* G233A in grey.

**Table 1.** Natural isolates tested for changes in mutation rate.

| Strain | Source | Sample | *cyp51A* | Clade | *pms1* | *msh2* | *msh6* |
| --- | --- | --- | --- | --- | --- | --- | --- |
| ARAF005 | Clinical | Sputum | Wild-type | B | Wild-type | Wild-type | Wild-type |
| C91 | Environmental | Plant Bulb | TR<sub>46</sub>/Y121F/T289A | B | Wild-type | Wild-type | Wild-type |
| C96 | Environmental | Plant Bulb | Wild-type | B | Wild-type | Wild-type | Wild-type |
| C136 | Clinical | Sputum | Wild-type | B | Wild-type | Wild-type | Wild-type |
| C272 | Environmental | Unknown | Wild-type | B | M315V | Wild-type | Wild-type |
| ARAF001 | Clinical | Sputum | TR<sub>34</sub>/L98H | A | E444G/S758Y | Wild-type | G233A |
| C1 | Environmental | Soil | TR<sub>34</sub>/L98H | A | Wild-type | Wild-type | G233A |
| C79 | Environmental | Compost | TR<sub>46</sub>/Y121F/T289A | A | Wild-type | K816Q | G233A/A1090S |
| C89 | Environmental | Plant Bulb | TR<sub>46</sub>/Y121F/T289A | A | Wild-type | E812G | Wild-type |
| C342 | Environmental | Soil | TR<sub>34</sub>/L98H | A | E444G | Wild-type | G233A |

## Discussion

The widespread use of agricultural azoles in the environment has selected for cross-resistance to critical clinical first line therapies used against *A. fumigatus* infections (*18*, *20*, *45*). As a result, a fungicide-driven multi-azole resistant clade has arisen typified by the expansion of the TR_34_/L98H allele in *cyp51A*. Our results show that isolates from this clade are not only resistant to azoles, but also have elevated mutation rates due to a linked variant allele of Msh6, a key protein in the MMR system involved in the recognition of base-base mispairings and insertion loops. This modest but significant increase in mutation rate in clade A isolates, increases the likelihood of resistance emerging to other antifungals in this clade. *A. fumigatus* produces vast numbers of spores and due to their mononucleate nature, even slightly elevated rates of mutation such as those measured here, mean it is highly likely that in any given niche where producing millions to billions of spores are produced, will have at least one spontaneous resistant mutant. Although mutational instability is associated with significant fitness costs in stable environments it is telling that deletion of *msh6* resulted in only a minor reduction in fitness costs *in vitro*. Though fitness costs may accumulate over longer evolutionary time periods, the relatively mild medium-term reduction in fitness that we observe might explain why the *msh6* G233A allele has persisted in *A. fumigatus* populations exposed to azole fungicides since their introduction in the mid-1970s (*46*, *47*). The complex lifecycle of *A. fumigatus* which can undergo both asexual and sexual reproduction (*48*) has also likely influenced the maintenance of the mutator phenotype (*49*). There are genetic signatures of high recombination between isolates within clade A, but there appears to be little recombination between clade A and B, which may aid in the maintenance of the mutator allele within clade A despite potential costs (*21*, *49*, *50*). Additionally, it would be expected that purifying selection would remove deleterious mutations from the population. This apparent reproductive isolation and purifying selection against costly mutations may help to explain the strong genetic linkage between clade A, TR_34_/L98H and the Msh6 mutator allele, its ongoing environmental persistence and would contribute to maintenance of the mutator phenotype within this clade.

Strong fluctuating selection through the extensive use of multiple different classes of fungicide may have provided strains with elevated mutation rates a fitness advantage by more rapid adaptation to changing abiotic stresses. Alternatively, as the Msh6 allele is closely linked to *cyp51A* on chromosome 4, it may be hitchhiking on strong directional selection for the TR_34_/L98H azole resistance mutation. Nevertheless, isolates from the multi-azole resistant clade A are between 4- and 9-fold more likely to acquire *de novo* mutations that provide resistance to novel antifungal therapies. Moreover, the impact of this elevation in mutation rate could be important in the within-patient evolution of antifungal resistance leading to more rapid treatment failure for CPA patients on long term antifungal therapy due to increased mutational supply. The advent of combination therapy through the use of novel, orally bioavailable classes of antifungal drugs with new modes of action including fosmanogepix (Phase II clinical trials) and olorofim (Phase III) should reduce the impact of antifungal resistance that arises in the patient. However, a number of agricultural antifungals have either been approved or are in late development for use in crop protection that share the same mechanism of action as these novel clinical drugs (*51*). The pyridine fungicide aminopyrifen (*52*) targets the same enzyme, GWT-1, as fosmanogepix. Likewise, ipflufenoquin which has been approved by the US Environmental Protection Agency for agricultural use (*53*) is analogous to the dihydroorotate dehydrogenase (DHODH) inhibitor olorofim. Further, ipflufenoquin has been shown to select for cross-resistance to olorofim (*54*). The dual use of these novel classes of antifungal drugs coupled with increased mutation rates in azole resistant isolates increases the probability of the accumulation of multiple independent resistance mechanisms within clade A to both azole and novel antifungal compounds. Through the pervasive anthropogenic use of agricultural azoles, a lineage of *A. fumigatus* has been selected for that is multi-azole resistant, multi-fungicide resistant, and also displays increased adaptability which ultimately will lead to the evolution of a lineage which manifests pan-drug resistance.

## Methods

### Strains, culture conditions and antifungals

*A. fumigatus* MFIG001 was used at the parental strain to generate *msh2*, *msh6*, and *pms1* deletion strains and *msh6* G233A mutant by CRISPR-Cas9 mediated transformation (*44*). *A. fumigatus* clinical and environmental isolates were provided by Matthew Fisher (*20*) and are described in Table 1. *A. fumigatus* strains used in this study were cultured on Aspergillus Complete Media (ACM) (*55*) agar for 3 days at 37°C unless stated otherwise. Spores were harvested in phosphate buffered saline (PBS) + 0.1% Tween-20 and filtered through miracloth (Millipore). Olorofim was synthesised by Concept Life Sciences, voriconazole was obtained from Merck (32483) and itraconazole from Cayman Chemical (13288). See Supplementary Table 2 for primers sequences used in this study.

### Gene Knockout and CRISPR-Cas9 gene modification

To construct the deletion mutants of MFIG001, mismatch repair genes *msh2*, *msh6* or *pms1* were replaced with the hygromycin resistance cassette. Target specific crRNAs and associated PAM sites flanking each gene were designed using EuPaGDT(*56*) based on the A1163 genome sequence. Homology directed repair (HDR) templates including the hygromycin resistance cassette were amplified from the pAN7.1 plasmid using primers that incorporated 50-bp microhomology arms flanking the gene to be replaced. CRISPR-Cas9 transformation was conducted as previously described (*44*). MFIG001 conidia were inoculated into liquid ACM and cultured for 16 hours at 37°C, shaking at 120 rpm. Mycelia were harvested by filtration through miracloth followed by proplasting using Vinotaste (Lamoth-Abiet) 10 g/mL in 0.6 M KCl + Citric acid at 37°C for 4 hours, shaking at 120 rpm. Protoplasts were harvested, centrifuged at 1,800 g for 10 mins, followed by washing three times with 0.6 M KCl. Protoplasts were resuspended in 0.6 M KCl + 200 mM CaCl_2_ and diluted to 1 × 10^6^ protoplasts/mL. RNP complexes were assembled using Alt-R *S.p.* Cas9 Nuclease V3, Alt-R® CRISPR-Cas9 tracrRNA and locus specific Alt-R® CRISPR crRNA (Integrated DNA Technologies) by heating to 95°C for 5 mins and gradually cooling to room temperature in a thermal cycler for 10 min. HDR template (1 µg), RNPs (33 µM RNA duplex with 1.5 µg Cas9) and protoplasts were mixed with PEG-CaCl_2_ (60% wt/vol PEG3350, 50 mM CaCl_2_, 450 mM Tris-HCl, pH7.5) and incubated on ice for 50 mins. Protoplasts were then spread onto YPS plates containing 100 μg/mL hygromycin (TOKU-E). Plates were incubated for 24 hours at room temperature followed by incubation at 37°C for 3 days. Transformants were purified and validated by PCR using primers flanking the loci (Supplementary Table 2), coupled with primers internal to the hygromycin resistance cassette, as well as primers specific to the loci deleted. For selection-tree CRISPR-Cas9 to recreate *msh6* G233A in MFIG001 a single stranded repair template was designed which incorporated the G698C transversion. Transformation was conducted as above and transformed protoplasts were plated onto non-selective YPS plates. Colonies were screened for *msh6* c.G698C by PCR using SNP specific primers (Supplementary Table 2), and positive transformants were confirmed by Sanger sequencing (Genewiz).

### Antifungal susceptibility testing

MIC determination for all drugs was carried out according to the broth microdilution method outlined by EUCAST(*57*). 1 × 10^5^ spores/ml were inoculated per well in CytoOne® 96-well plates (StarLab) containing RPMI-1640 medium supplemented with 2.0% glucose and MOPS pH 7.0. Growth was measured by OD at 600 nm in a two-fold dilution series of antifungal drug (olorofim 0.00195-2 µg/ml, voriconazole 0. 0156-16 µg/ml) following incubation at 37 °C for 48 h. Three biological replicates for each strain tested were conducted. The MIC was defined as the lowest concentration required to reduce growth by 90%.

### Fluctuation test and mutation rate analysis

Fluctuation tests were performed by inoculating 12 independent 10 ml cultures with 5000 spores on ACM agar from a single spore stock and allowed to grow in non-selective conditions for 72 hours at 37°C. To determine the effect of the mutagen ethyl methanesulfonate on mutation rate of MFIG001, a two-fold series of EMS, between 64 and 512 µg/ml, was added to the growth media. Spores were harvested in PBS + 0.1% tween-20, spores were then pelleted by centrifugation and resuspended in 500 µl PBS + 0.1% tween-20. 2 µl of spores from each culture were removed, diluted, and used to measure the final population size (*N_t_*) by flow cytometry. We obtained the number of spontaneous resistant mutants that formed during growth in non-selective conditions by transferring the total remaining spores to ACM agar plates containing either 2 µg/ml voriconazole, 8 µg/ml itraconazole or 0.5 µg/ml olorofim. Plates were incubated for 72 hours at 37°C. For *Δmsh2*, *Δmsh6* and *Δpms1* (Figure 2b) and *msh6* G233A (Figure 4a/b) mutation rates against voriconazole and olorofim were calculated. An estimate of the number of mutation events (*m*) was calculated from the distribution of the number of mutants across the 12 independent cultures using the Ma-Sandri-Sarkar maximum-likelihood method implemented in the *flan* R package. To calculate the mutation rate per spore *m* was divided by the final population size *N_t_*. For *Δmsh2*, *Δmsh6* and *Δpms1* mutation frequency to itraconazole and olorofim (Figure 2c,d) were calculated from 6 independent cultures by dividing the number of resistant colonies by the number of spores plated. For each fluctuation test at least 3 randomly selected spontaneous resistant mutants were sub-cultured on non-selective ACM for a further 72 hours at 37°C followed by culturing on ACM containing 2 µg/ml voriconazole, 8 µg/ml itraconazole or 0.5 µg/ml olorofim to confirm resistance. The genes encoding targets of voriconazole, *cyp51A,* and olorofim, *pyrE,* of 27 voriconazole and 27 olorofim spontaneous resistance mutants were also Sanger sequenced.

### Fitness assays

Radial growth rates of *Δmsh2*, *Δmsh6* and *Δpms1* mutants were conducted by spotting 10^3^ spores per strain on ACM or Aspergillus Minimal Media (AMM) agar plates. Plates were incubated for 96 hours and the radius of the colonies were measured every 24 hours. Three biological replicates were conducted for each strain and media condition. To measure fitness over longer time scales, culture flasks containing 10 ml of ACM or AMM were inoculated with 1:1 mixture of MFIG001 plus either *Δmsh2*, *Δmsh6* and *Δpms1* mutant strains at a final density of 5000 spores/ml. The initial ratio of WT to mutant was determined by diluting the spore mixture to appropriate concentration and plating onto non-selective and hygromycin plates to calculate the total population size and fraction of KO mutant in the mixture. The cultures were incubated for 72 hours at 37°C at which point total spores were harvested, 1% of the total harvested spores were used to inoculate fresh ACM or AMM. The remaining was diluted and plated on non-selective and hygromycin plates to calculate the frequency of deletion mutant within the population. The was repeated for a total of 5 transfers.

### Bioinformatics

Analysis of variants in MMR genes was conducted on 218 previously sequenced *A. fumigatus* isolates (ENA:PRJEB27135)(*20*). Orthologous sequences of mismatch repair genes *msh2* (AFUB_039320, AFUA_3G09850), *msh3* (AFUB_090020, AFUA_7G04480), *msh6* (AFUB_065410, AFUA_4G08300), *pms1* (AFUB_029050, AFUA_2G13410) and *mlh1* (AFUB_059270, AFUA_5G11700) were extracted from the previously published pan-genome(*20*) by identifying the representative pan-genes using BLASTP v2.12.0. Extracted sequences underwent multiple-sequence alignment using MUSCLE v3.8.1551(Ref. (*58*)) and variants were identified using SNP-sites v2.5.1 (Ref. (*59*)) and confirmed by visualization in JalView v2.11.2.6 (Ref. (*60*)). Whole-genomic single nucleotide polymorphism (SNP) phylogeny of UK isolates was produced as described in Rhodes *et al*(*20*). Phylogenetic trees with overlaying metadata were generated using iTOL v6.5.4 (Ref. (*61*)).

For analysis of global isolates publicly available raw reads (Table S2) were mapped to *Af*293 (GCF_000002655.1) using Burrows-Wheeler Aligner v0.7.17 (Ref. (*62*)). Text pileup outputs were generated for each sequence using SAMtools v1.6 mpileup with option –I to exclude insertions and deletions (*63*). BCFtools v1.6 call was used to call SNPs with options -c to use the original calling method and --ploidy 1 for haploid data (*64*). Consensus genome sequences in fasta format were extracted from vcf files using seqtk v1.2 (available at https://github.com/lh3/seqtk[github.com]) with bases with a phred quality score below 40 counted as missing data. MEGA version X (*65*) was used to create a neighbour-joining tree using the Tamura-Nei model with 100 bootstraps for all whole genome sequences. The phylogeny was visualized using iTOL (*61*).

Sequence conservation at the G233 locus of *msh6* was examined in 289 members of the family Trichocomaceae retrieved from the NCBI non-redundant protein database through protein blast using *Af*293 Msh6 (AFUA_4G08300) as the reference. Sequences were aligned using MUSCLE v3.8.1551 and sequence logos were created using WebLogo (*66*).

### Statistics

All statistics were conducted in R v4.1.1. Assumptions of normality were determined using Q-Q plots and Shapiro-Wilk tests, and where appropriate non-parametric tests were conducted. Pearson’s chi-squared tests were used to test for significant difference between expected and observed frequencies of variant alleles between clade A and clade B, and between TR_34_/L98H and WT *cyp51A* genotypes. Significance differences between mutation rates were calculated using maximum likelihood estimation using the Luria-Delbrück model (*flan* package R). Kruskal-Wallis test tested for significant differences in MIC of spontaneous resistant mutants in *Δmsh2*, *Δmsh6*, and *Δpms1* backgrounds. Pair wise differences between groups were determined by ANOVA with Tukey post hoc tests where assumptions were met and Wilcoxon rank sum tests otherwise. To model the effect of transfer, strain and media upon fitness a mixed effects linear model was fitted to the data using the *nlme* package in R. The model fit the frequency of mutant strain to the interacting effects of transfer, strain, and media type, with replication modelled as a random effect. The model allowed variance to change with fitted value to account for the reduced variance as mutant frequency approached zero.

