## Supplemental Figures S1-S5 for "Elevated mutation rates in the multi-azole resistant *Aspergillus fumigatus* clade drives rapid evolution of antifungal resistance"

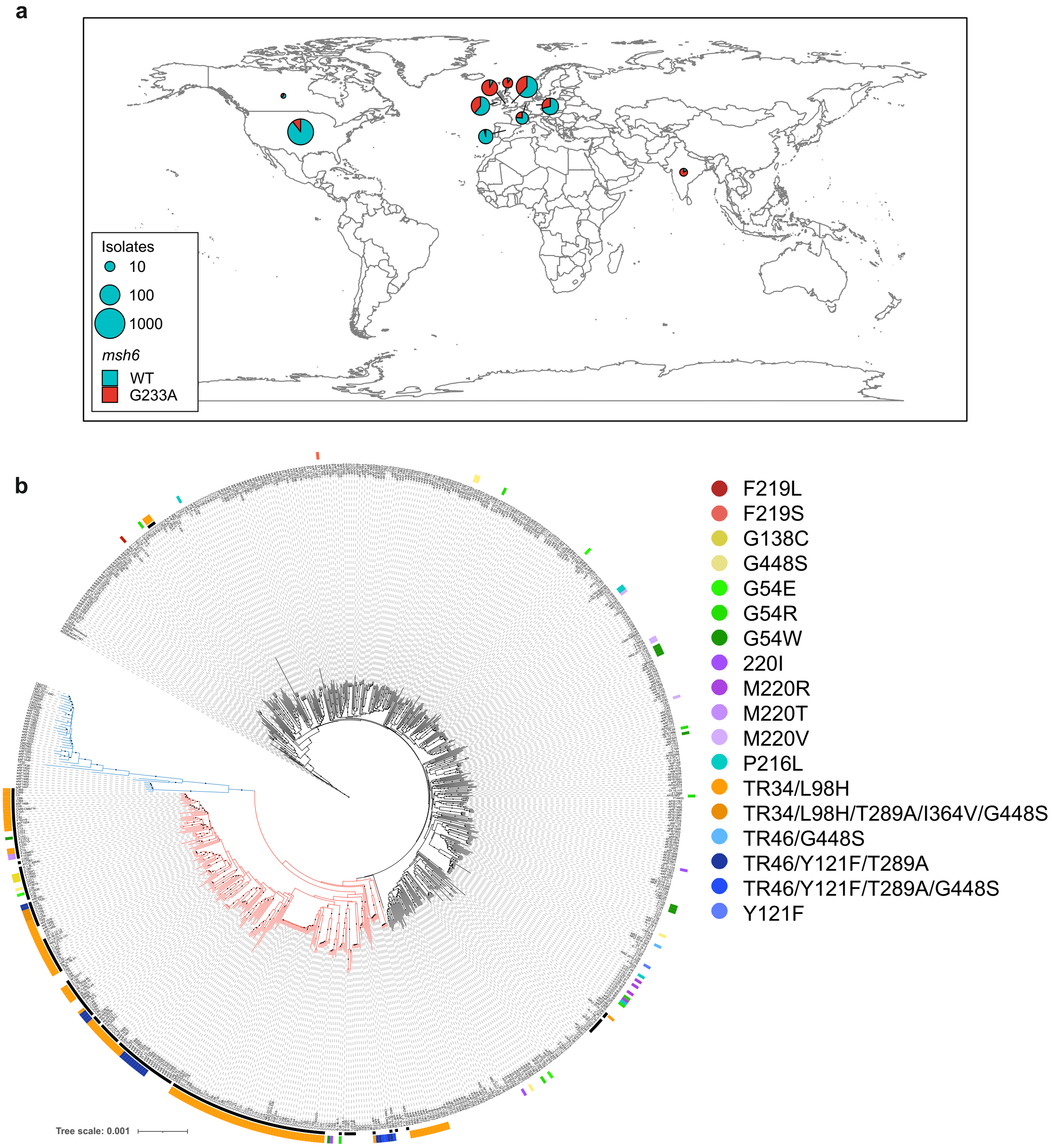


**Figure S1 Global presence of G233A within $TR_{34}$/L98H genotypes. a** Map showing the proportion of *msh6* G233A variants by country for 728 publicly available whole genome sequenced *A. fumigatus* isolates show that the mismatch repair variant is observed globally. Size of pie charts scaled by number of isolates. **b** A neighbour joining phylogenetic tree using genome wide SNPs supports the clustering of $TR_{34}$/L98H genotypes in clade A (red branches) and the association with *msh6* G233A variant (black points on tip labels) rooted to Af293.

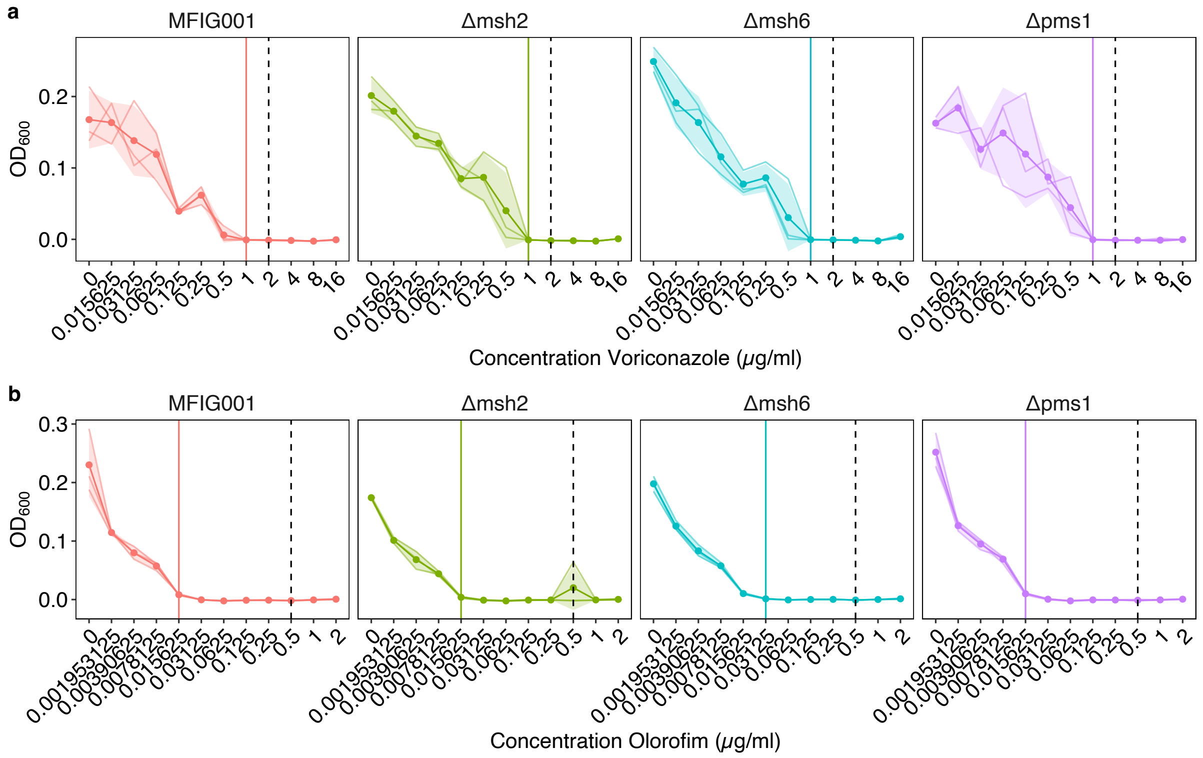


**Figure S2 MMR deficient mutants do not show altered MIC to voriconazole or olorofim. a** MIC curves of MFIG001 parental strain and *Δmsh2*, *Δmsh6* and *Δpms1* to voriconazole. **b** MIC curves of MFIG001 parental strain and *Δmsh2*, *Δmsh6* and *Δpms1* to olorofim. Bold lines show means of three replicates, each of which are shown separately. Shaded area shows +/- standard deviation. Dashed vertical line shows concentration used to select for resistant mutants, vertical solid line shows MIC.

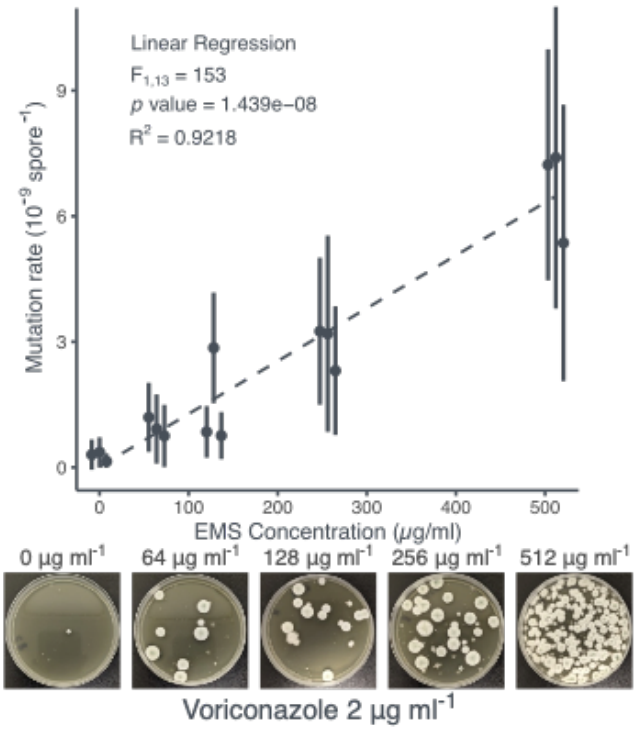


**Figure S3 the effect of ethyl methanesulfonate on mutation rate.** Mutation rates for resistance to voriconazole for the MFIG001 reference strain. Each point shows the calculated mutation rate from a single independent fluctuation test using 12 replicate cultures. Error bars show 95% confidence intervals. Dashed line shows linear regression fit. Images of example selective plates shown below plot.

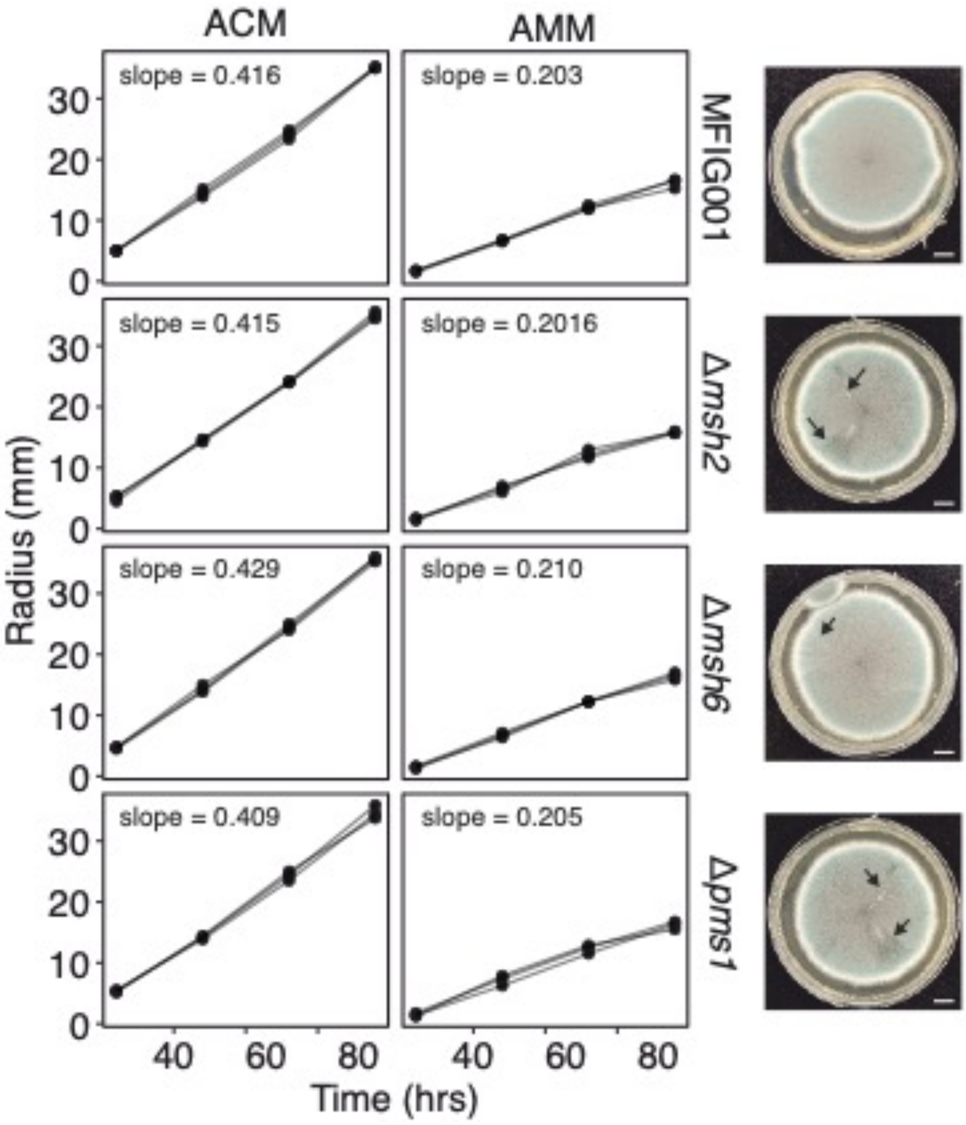


**Figure S4 Radial growth rates of MMR deficient mutants.** Radial growth rate of MFIG001 parental strain and *Δmsh2*, *Δmsh6* and *Δpms1* mutants in aspergillus complete media (ACM) and aspergillus minimal media (AMM), N = 3. Images of growth after 96 hours, arrows highlight presence of sectoring. Scale bar represents 10 mm.

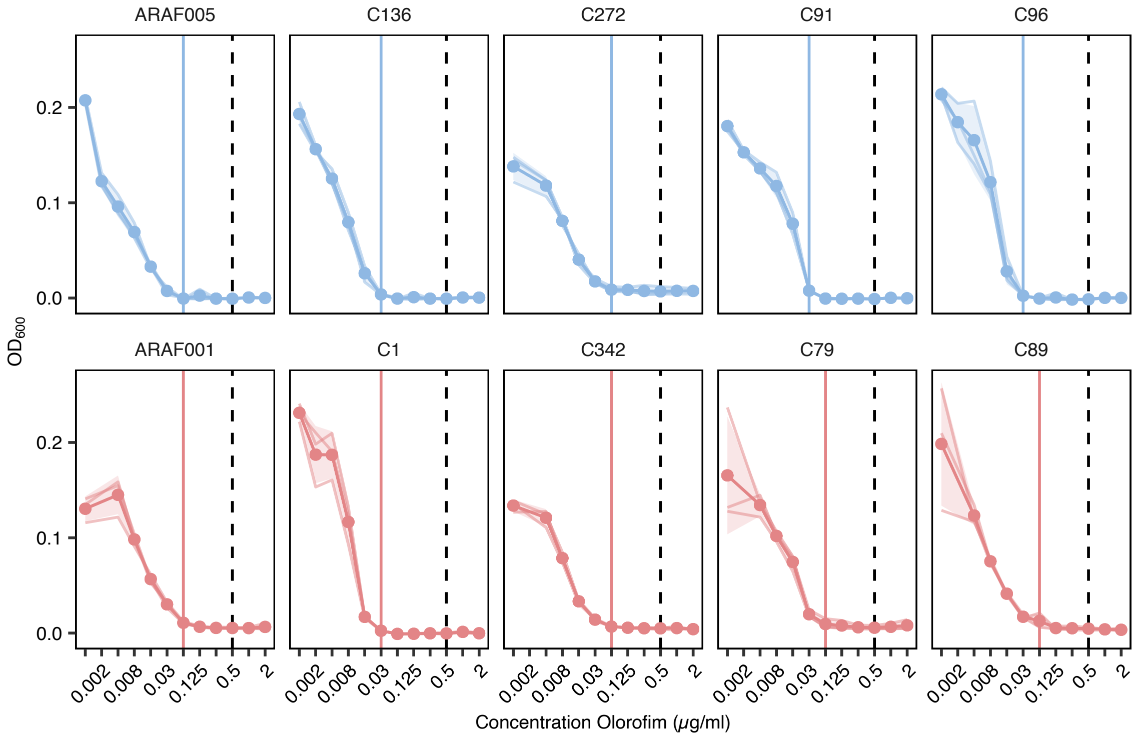


**Figure S5 MIC curves of natural isolates to olorofim.** MIC curves of 10 natural isolates with olorofim. Bold lines show means of three replicates, each of which are shown separately. Shaded area shows +/- standard deviation, N = 3. Dashed vertical line shows concentration used to select for resistant mutants, vertical solid line shows MIC. Coloured by clade, red clade A, blue clade B.
